# Oral Administration of a specific p300/CBP Lysine Acetyltransferase Activator Induces Synaptic Plasticity and Repairing Spinal Cord Injury

**DOI:** 10.1101/2023.10.04.560982

**Authors:** Akash Kumar Singh, Amrish Rai, Ila Joshi, Damodara N Reddy, Rajdeep Guha, Shubha Shukla, Aamir Nazir, James P. Clement, Tapas K Kundu

**Author notes:** Equal contribution (Akash Kumar Singh, Amrish Rai).

## Abstract

TTK21 is a small molecule activator of p300/CBP acetyltransferase activity, which upon conjugation with glucose-derived carbon nanosphere (CSP) can efficiently cross the blood-brain barrier and activate histone acetylation in the brain. Its role in adult neurogenesis and retention of long-term spatial memory upon intraperitoneal (IP) administration is well established. In this study, we have successfully demonstrated that CSP-TTK21 can be effectively administered via oral gavage. Using a combination of molecular biology, microscopy, and electrophysiological techniques, we systematically investigated the comparative efficacy of oral administration of CSP and CSP-TTK21 in wild-type mice, evaluating their functional effects in comparison to intraperitoneal (IP) administration. Our findings indicate that CSP-TTK21, when administered orally, induces long-term potentiation in the hippocampus without significantly altering basal synaptic transmission, a response comparable to that achieved through IP injection. Remarkably, in a spinal cord injury model, oral administration of CSP-TTK21 exhibits equivalent efficacy to IP administration. Furthermore, our research demonstrates that oral delivery of CSP-TTK21 leads to improvements in motor functions, histone acetylation dynamics, and increased expression of regeneration associated genes (RAGs) in a spinal injury rat model, mirroring the effectiveness of IP administration. Collectively, these results underscore the potential utility of CSP as an oral drug delivery system, particularly for targeting the neural system.

## INTRODUCTION

Memory, the capacity to encode, retain, and retrieve information as needed for daily activities, relies on intricate processes involving dynamic alterations in gene expression. These changes reshape the structure and function of neuronal synapses. Depending on whether information is stored, recalled, or erased, dendritic spines and neuronal synapses undergo substantial modulation and are termed synaptic plasticity [1, 2]. However, the molecular mechanism driving synaptic plasticity and its impact on memory and cognition remains largely uncharted. Epigenetic changes, specifically DNA methylation and histone acetylation, play pivotal roles in memory and cognitive processes [3, 4]. Utilizing lysine deacetylases (KDAC) inhibitors (KDACi) to boost histone acetylation has demonstrated the potential to improve learning and cognitive abilities in mouse models [5, 6]. Moreover, recent research highlights the therapeutic promise of KDACi in treating various neurodegenerative disorders [7–10]. Due to their pleiotropic effect and limited specificity, these molecules could not proceed further for clinical trials [11, 12]. Thus, an alternative use of KDACi is lysine acetyltransferases (KAT) activators. One such molecule is TTK21, a novel KAT activator that specifically activates p300/CBP-KAT activity *in vitro* as well as *in vivo* [13–16]. Previous research in our laboratory demonstrated that when administered via intraperitoneal injections in mice, the CSP-TTK21 molecule, coupled with glucose-derived carbon nanospheres (CSP), could cross the blood-brain barrier (BBB) and induce histone acetylation in the dorsal hippocampus and frontal cortex. Furthermore, it facilitated adult neurogenesis and significantly extended long-term spatial memory retention [13]. In this study, we explored the feasibility of orally administering CSP-TTK21, a more common and user-friendly route for humans. We also conducted bio-distribution studies and compared the activity of CSP-TTK21 between the two administration methods: intraperitoneal (IP) and oral gavaging.

CBP/p300 lysine acetyltransferases are the master regulators of gene expression and are known to induce regenerative growth of DRG neurons [17, 18]. Previously, we have demonstrated that by pharmacologically activating CBP via intraperitoneal administration of CSP-TTK21, induced axon regeneration, sprouting, and facilitated functional recovery in both acute and chronic rodent models of spinal cord injury (SCI) [15, 19]. CSP-TTK21 was shown to enhance neuronal activity by epigenetic modifications, in particular inducing CBP-mediated histone acetylation, which increased expression of regeneration-associated genes leading to increased regenerative potential and functional recovery after SCI. However, there is still a gap in understanding the pharmacological activation of CBP through oral delivery of CSP-TTK21 in non-clinical SCI models, which is crucial for bridging the divide between preclinical and clinical research. Here, we show that oral delivery of CSP-TTK21 restored acetylation dynamics and enhanced motor functions in a rat model of spinal cord injury. Mechanistically, we found that CSP-TTK21 treatment increased expression of RAGs in different brain regions such as prefrontal cortex and cerebellum which are associated with altered structural and functional activity after spinal cord injury.

## RESULTS AND DISCUSSION

### CSP-TTK21 exhibits BBB penetration upon oral administration and remains active

For this study, CSP and CSP-TTK21 were synthesized, conjugated, and characterized (Fig. 1) following procedures similar to those detailed in previously [13, 20]. The identical batch was consistently employed throughout the study. We began with a pilot experiment to assess the viability of orally delivering this compound. Through immunohistochemistry (IHC) of mice that had been fed CSP or CSP-TTK21, we successfully detected CSP in the hippocampus of these mice (Fig. 2, A and B). Moreover, the conjugated TTK21 exhibited activity, as evidenced by the induction of the H4K12ac mark in the dorsal hippocampus. (Fig. 2C). We observed that the highest number of nanospheres were present in the brain on day 4 after oral gavaging, in contrast to day 3, as previously seen with IP administration.

**Fig. 1.**
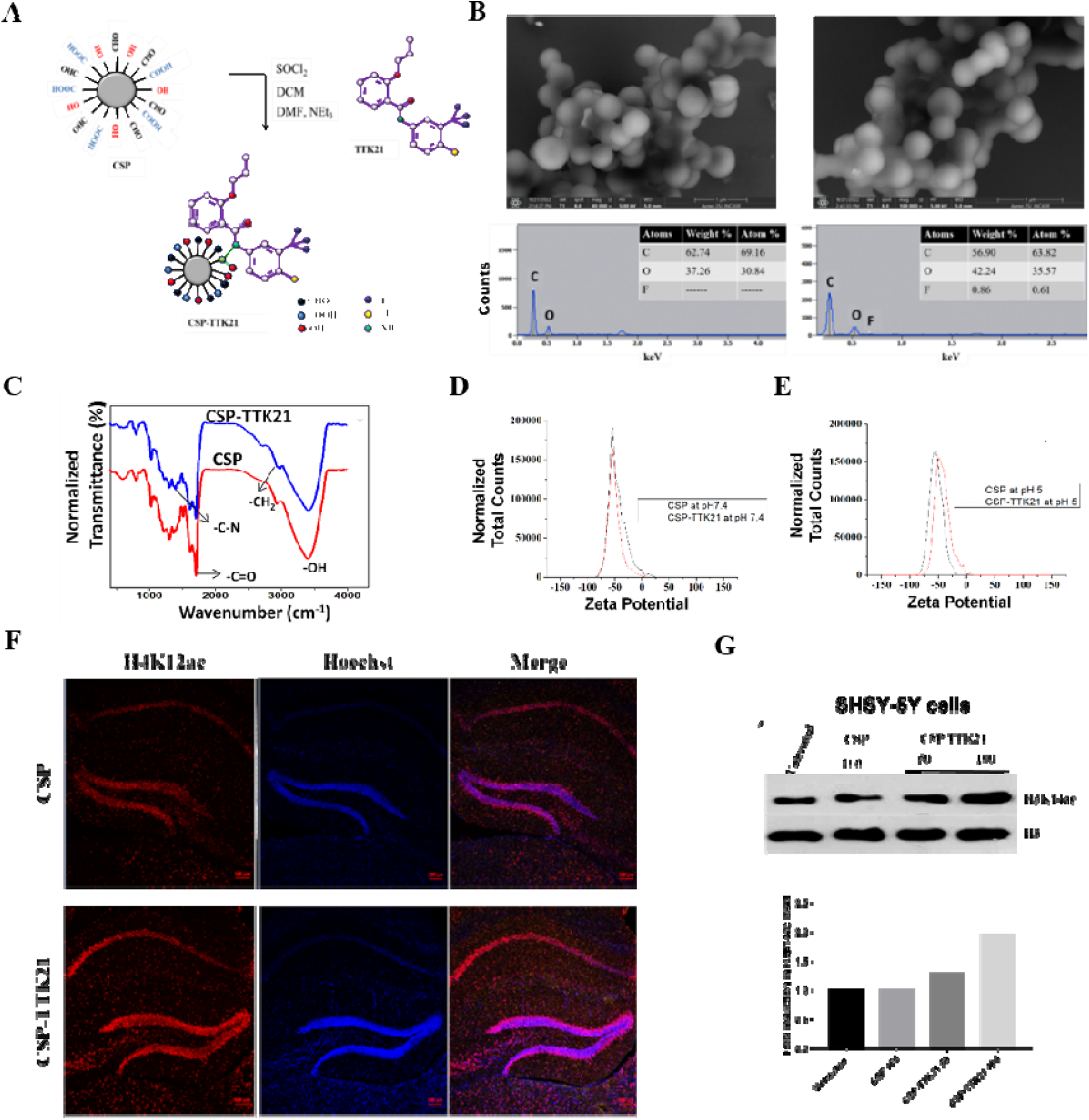
Conjugation and CSP-TTK21 characterization. (A) Schematic representation of CSP and TTK21 conjugation. (B) FESEM images of CSP and CSP-TTK21, presence of F-atom confirms CSP-TTK21 conjugation. (C) NMR spectra of CSP and CSP-TTK21. (D) Zeta potential of CSP and CSP-TTK21 at pH 7.4 and (E) at pH 5. (F) Immunofluorescence image showing increased H4K12ac marks in CSP-TTK21 treated mice hippocampus as compared to CSP mice. (G) Dose dependent increase in H3K14ac marks upon CSP-TTK21 treatment in SHSY5Y neuroblastoma cells. Scale bar (B) 1 µm and (F) 100 µm.

The reduced presence of fluorescence CSP particles in the brain following oral gavaging, compared to IP administration, can be attributed to the fact that IP injections deliver molecules directly into the peritoneal cavity, ensuring efficient absorption into the bloodstream, which is relatively quicker. Conversely, during oral delivery, the molecules must traverse the esophagus, stomach, and gastrointestinal (GI) tract before reaching the bloodstream, a process that takes more time. Despite the presence of proteases and digestive enzymes in the stomach and GI tract, our observation that CSP-TTK21 actively induces histone acetylation in the brain, especially in the hippocampal region, suggests that the molecules remain intact after this entire journey (Fig. 2, D-F). However, we cannot rule out the possibility of some degree of CSP particles being digested during oral gavaging due to their glucose-based carbon nanosphere composition. Since our primary goal was to evaluate the functionality of these molecular conjugates in the brain following oral delivery, these results provide valuable insights into the potential viability of CSP-TTK21 being delivered orally and retaining its expected functionality.

**Fig. 2.**
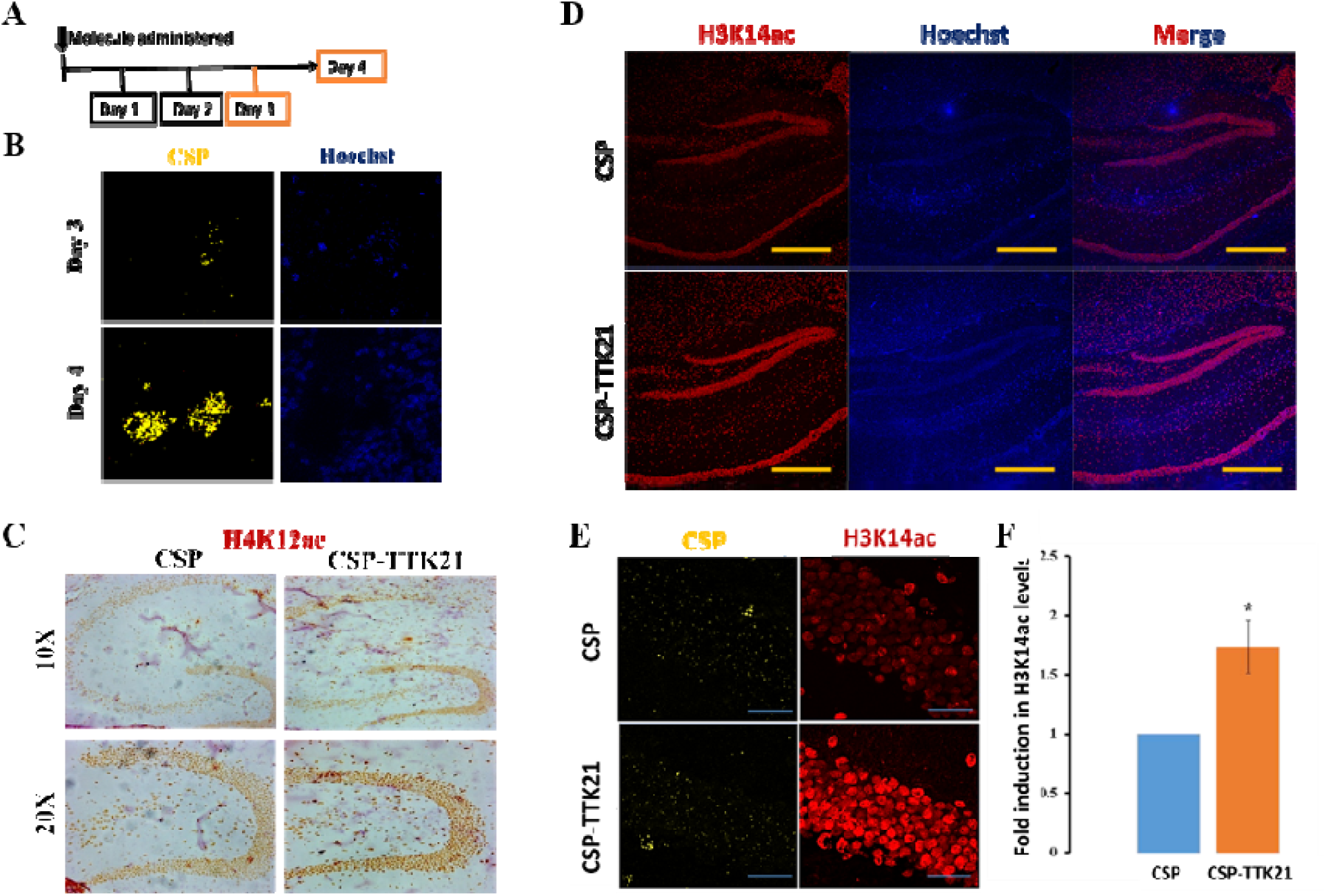
Oral administration of CSP-TTK21. (A) Schematic showing experimental timeline. (B) Presence of CSP particles in brain at different days (2 mice/group). (C) DAB staining showing induction of H4K12ac marks in CSP-TTK21 treated hippocampal region (2 mice/group). (D) Immunostaining showing induction of H3K14ac marks in the Hippocampus (5-6 mice/group. (E) Enlarged image at 63X magnification showing the presence of CSP particles as well as induction of H3K14ac marks upon CSP-TTK21 treatment. (F) Quantification of fold induction in H3K14ac levels in CSP-TTK21 treated condition as compared to CSP control. Scale bar (d) 200 µm and (e) 20 µm.

### Bio-distribution of CSP following IP and Oral administration

We investigated CSP biodistribution upon oral gavaging and studied its patterns across various mouse organs. Mice received a single CSP dose (∼20 mg/kg) through oral and IP administration. We analyzed samples from 6 hours to 21 days post-treatment (Fig. 3A). The highest particle concentration in the hippocampus occurred on day 1 in both administration modes, declining by day 3 and nearly vanishing by day 21 (Fig. 3, B and C). With oral administration, we observed initial particle accumulation in the liver on day 1, decreasing by day 3 and almost disappearing by day 21. A similar, albeit less pronounced, pattern was seen with IP administration, diminishing from day 3 to undetectable levels by day 21. Kidney showed early particle presence on day 1 (oral) and day 3 (IP), reducing by day 21.

**Fig. 3.**
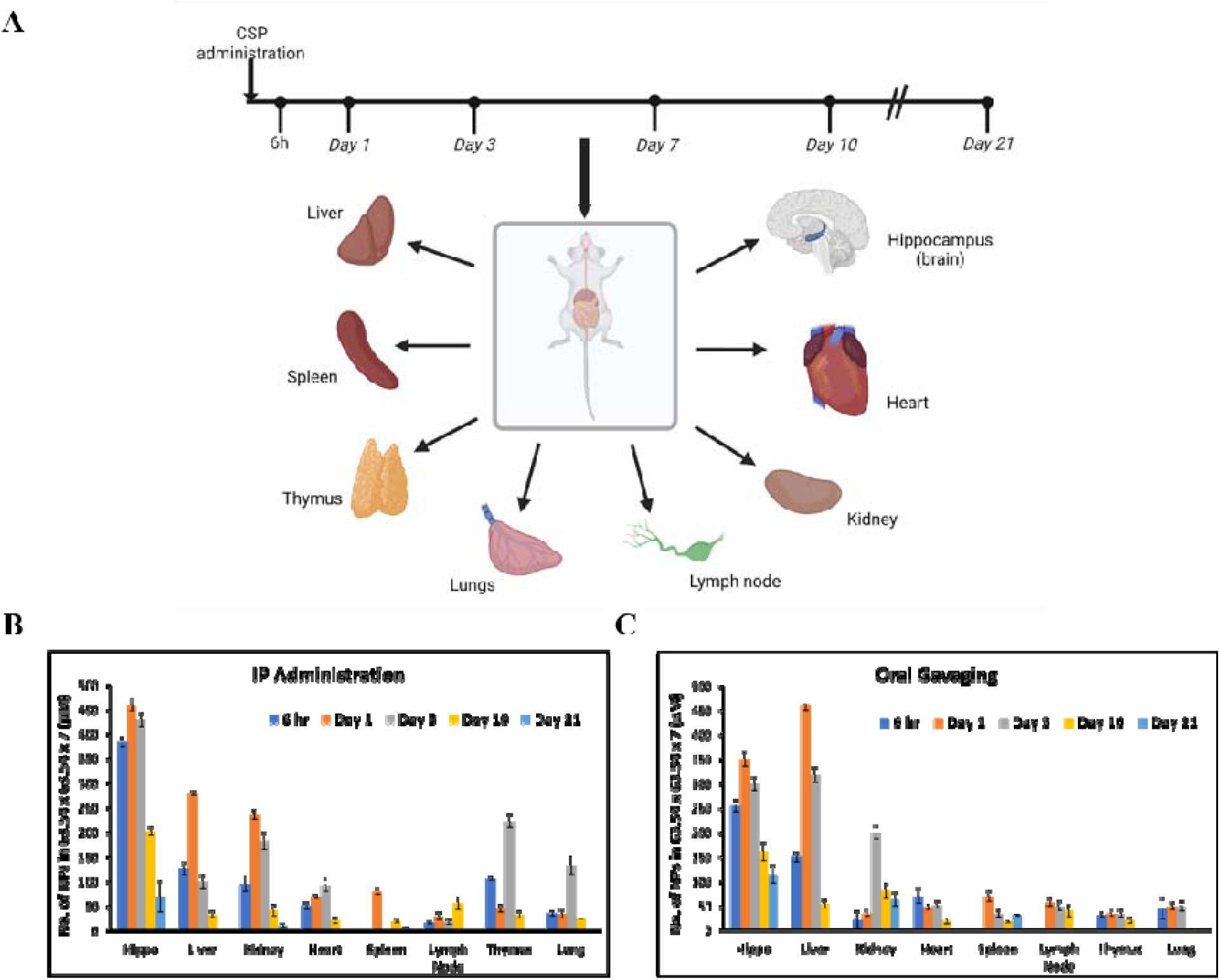
Bio-distribution study of CSP upon IP and Oral administration by confocal imaging. Mice were treated with single dose of CSP and sacrificed at different time intervals (3 mice/group). (A) Schematic showing the experimental timeline and the organs used for bio-distribution studies (Created with BioRender.com). (B) Bio-distribution profile of CSP in various organs upon IP administration. (C) Bio-distribution profile of CSP in various organs upon oral administration.

Additionally, particles were found in the heart, spleen, lymph node, thymus, and lung in both routes but at lower levels, clearing by day 21 (Fig. 3). Importantly, no changes in tissue morphology were observed during these studies. These findings indicate CSP’s favorable bioavailability, pharmacokinetics, and apparent lack of toxicity. It suggests CSP’s potential use as a drug carrier for conditions like obesity, fatty liver disease in the liver, and cystinosis, along with other acute/chronic kidney diseases.

### Oral Gavaging is equally effective as IP injections

We conducted a comparative analysis to assess the effectiveness of CSP-TTK21 through oral delivery in comparison to IP injection. We evaluated the activation of p300 lysine acetyltransferase (KAT) activity for both administration routes. Since we observed the highest particle presence at day 4 instead of day 3, which was typically seen with IP injections, we opted for a middle time point at 84 hours (3 ½ days) for comparison purposes. Two groups of mice received a single dose of CSP and CSP-TTK21: one through oral gavaging and the other via IP injections. After 84 hours, the mice were euthanized, and each hemisphere of the brain underwent confocal imaging and electrophysiological studies. Immunohistochemistry revealed a noteworthy increase in histone H3K14 acetylation levels in the dorsal hippocampus of mice treated with CSP-TTK21, regardless of the administration route (Fig. 4, A and B). Furthermore, a comparison of acetylation activation between the two conditions (oral delivery vs. IP injection) showed no significant difference, indicating that CSP-TTK21’s oral delivery is equally effective as IP injections.

The specific modification (acetylation) of lysine14 residue on histone H3 has previously been reported to occur during activation of some neuronal receptors such as dopamine receptor, muscarinic-acetylcholine receptor, and glutamate receptor [21]. Subsequently, we assessed the impact of CSP-TTK21 on basal synaptic transmission and long-term potentiation (LTP) in these mice through extracellular field recordings in the hippocampal region (Schaffer collateral pathway). We observed that CSP-TTK21 treatment did not modify basal synaptic transmission compared to the CSP control, as evident from the input-output curve (Fig. 4C). This indicates that the molecule had no discernible effect on basal synaptic transmission. Further, we studied the effect of this compound on LTP, by conducting extracellular field recordings from the Schaffer collateral pathway in the hippocampus. Our findings revealed that this molecule markedly enhanced long-term potentiation in mice treated with a single dose of CSP-TTK21 compared to those treated with CSP alone. Intriguingly, when we compared the extent of LTP induction in both scenarios (oral and IP administration), we observed a remarkably similar magnitude of LTP enhancement in both cases (Fig. 4, D and E). This outcome strongly validates our earlier observation that the oral delivery of CSP-TTK21 is as effective as IP injections.

**Fig. 4.**
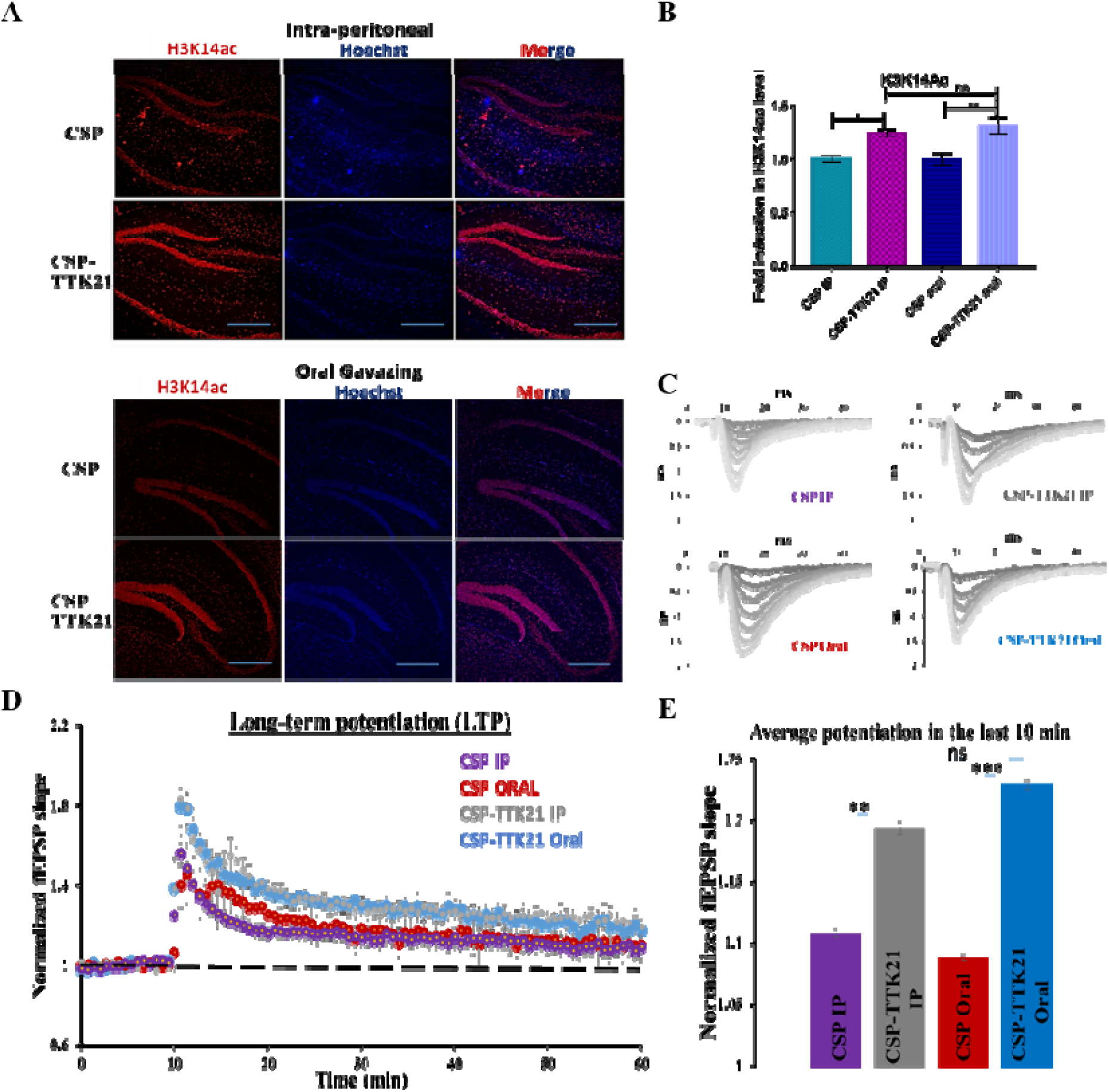
Comparison of CSP-TTK21 activity upon IP and Oral administration. (A) Immunohistochemistry was performed on 20µm thick sections with an anti-acetylated H3K14 antibody. In both cases IP and Oral administration, acetylation was increased in the hippocampal region as compared to CSP-control (n=4). Scale bar 200µm. (B) Quantitation of fold induction in histone acetylation level upon Oral and IP administration of CSP-TTK21 (normalized to CSP). (C) Sample traces for input-output curve analysis for each group. (D) Induction of LTP upon CSP-TTK21 treatment both with oral and IP injection. (E) Quantitation showing a significant increase in potentiation during the last 10 min of recordings in CSP-TTK21 treated group. Error bars represent the standard error of the mean (SEM). Student’s t-test was done for significance test, *p<0.05.

### CSP-TTK21 promotes motor recovery after spinal cord injury

Sensory-motor functions gradually fail because of aberrant activity in the spinal circuits below the site of injury [22]. Accumulative evidence suggests that epigenetic modifications such as acetylation can support the regeneration in a transcription dependent manner [23, 24]. We and others have shown that histone lysine acetyltransferase CBP/p300, is an essential epigenetic factor, which acetylates and induces several regeneration-associated growth signals in DRG neurons [15, 17]. Importantly, we found that systemic injection of small molecule activator (CSP-TTK21) of CBP/p300 promotes axonal regeneration and function recovery after spinal cord injury (SCI) [15]. Here, we investigated the effects of oral delivery of CBP/p300 KAT activator CSP-TTK21 in a rat SCI model. To test whether oral delivery of CSP-TTK21 can promote functional recovery, we have utilized a mid-thoracic dorsal hemisection model of spinal cord injury as previously reported [15, 19]. Briefly, the SCI model was generated as shown in (Fig. 5, B and C). CSP-TTK21 was administered 4 hours after the spinal injury and treatment was continued until the conclusion of the behavioral experiments with one dosage per week (Fig. 5A). We observed SCI rats that were treated with CSP-TTK21 showed improved locomotion activity in 1st week (Fig. 5, D and E) and by 5^th^ week there was significant enhancement in locomotion and rearing activity in these rats (Fig 5, F and G). These results suggest that oral delivery of CSP-TTK21 executed a similar effect as systemic injection, promoting functional recovery after SCI.

**Fig. 5.**
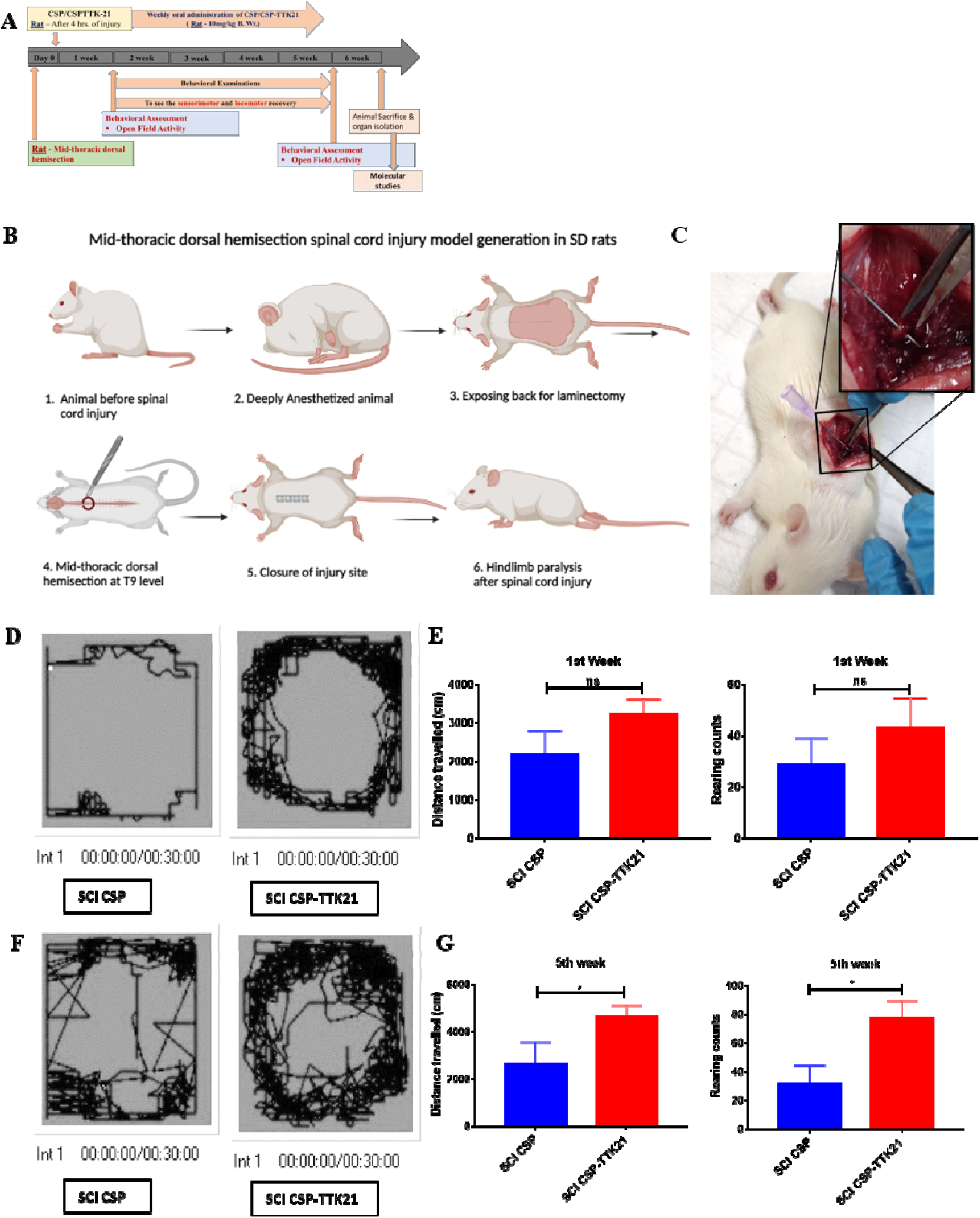
Locomotor behavioral test in CSP-TTK21 treated rats in spinal cord injury. (A) Schematic showing experimental timeline. (B) Schematic showing SCI model generation in SD rats (Created with BioRender.com). (C) Mid-thoracic dorsal hemisection at T9. (D) Representative tracks in OFT and (E) Total distance travelled and rearing counts during the 30 min OFT session after 1st week of CSP-TTK21 treatment (5 mice/group). (F) Representative tracks in OFT and (G) Total distance travelled and rearing counts during the 30 min OFT session after 5th week of CSP-TTK21 treatment (5 mice/group). Student’s t-test was done for significance test, *p<0.05.

### CSP-TTK21 induces histone acetylation in spinal cord after spinal cord injury

Motor function recovery after SCI has been associated with epigenetic modifications leading to expression of regeneration-associated genes (RAGs). Previous findings suggest histone acetylation dynamics as one of the critical factors in expression of the RAGs and axonal regeneration after SCI. Therefore, we next investigated the effect of oral delivery of CSP-TTK21 on histone acetylation dynamics in SCI rats. By immunofluorescence analysis, we found that H4K12ac was enhanced in the dorsal horn of spinal cord in SCI rats treated with CSP-TTK21 as compared to CSP vehicle rats (Fig. 6 A-C). Interestingly, another acetylation mark, H3K27ac that is associated with enhancer region was also observed to be significantly increased in the dorsal horn of spinal cord in CSP-TTK21 treated SCI rats (Fig. 6 D-F). To further strengthen our evidence, we next performed immunoblotting from spinal cord samples of these rats. We observed increased H3K9ac levels in spinal cord of SCI rats treated with CSP-TTK21 as compared to CSP vehicle control (Fig. 6 G and H). Together these results suggest that CSP-TTK21 via activation of CBP/p300 activity induces histone acetylation in dorsal horn of spinal cord which may lead to expression of RAGs.

**Fig. 6.**
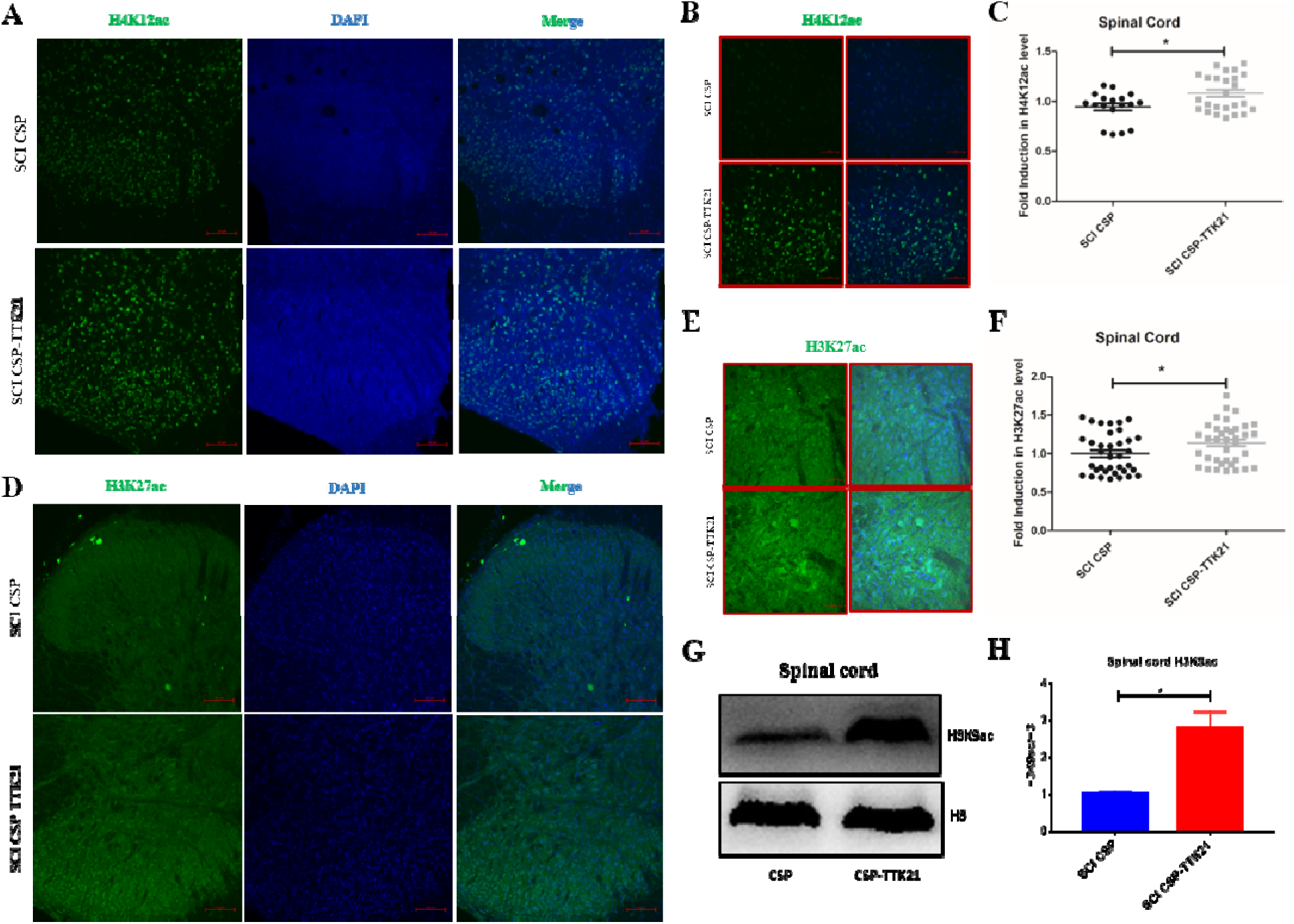
Histone acetylation in dorsal horn of spinal cord in CSP-TTK21 treated rats in spinal cord injury. (A) Immunofluorescence image showing increased H4K12ac (green) in dorsal horn of spinal cord. (B) Enlarged image at 40X magnification showing the induction of H4K12ac marks upon CSP-TTK21 treatment. Scale bar (A) 100 µm and (B) 50 µm. (C) Quantification of fold induction in H4K12ac levels in dorsal horn of spinal cord of SCI CSP-TTK21 rats as compared to SCI CSP rats (4-5 rats/group). (D) Immunofluorescence image showing increased H3K27ac (green) in dorsal horn of spinal cord. (E) Enlarged image at 40X magnification showing the induction of H3K27ac marks upon CSP-TTK21 treatment. Scale bar (D) 100 µm and (E) 50 µm. (F) Quantification of fold induction in H3K27ac levels in dorsal horn of spinal cord of SCI CSP-TTK21 rats as compared to SCI CSP rats (4-5 rats/group). (G) Immunoblot analysis showing increased H3K9ac marks in spinal cord (H) Quantification of fold induction in H3K9ac in spinal cord of SCI CSP-TTK21 rats as compared to SCI CSP rats (3-4 rats/group). Student’s t-test was done for significance test, *p<0.05.

### CSP-TTK21 induces histone acetylation in prefrontal cortex and cerebellum after spinal cord injury

Spinal cord injury result in a global structural and functional changes in different brain regions such as medial prefrontal cortex (mPFC), primary motor cortex, somatosensory cortex and cerebellum leading to reduced grey matter volume [25–27]. However, changes in histone acetylation majorly in prefrontal cortex and cerebellum with its role in regulating motor activity and regeneration in spinal injury is not much explored. To gain deeper insights into the role of CBP/p300 mediated induction of histone acetylation and regeneration after spinal cord injury, we examined the effect of orally administering CSP-TTK21 on histone acetylation dynamics in prefrontal cortex and cerebellum. By immunofluorescence analysis, we observed increased level of H4K12ac marks in prefrontal cortex of SCI rats treated with CSP-TTK21 as compared to vehicle control SCI rats (Fig. 7, A-C). A similar effect was visible for H3K27ac (Fig. 7, D-F). Moreover, immunoblotting also showed increased H3K9ac levels in the prefrontal cortex of CSP-TTK21 treated SCI rats (Fig. 7, G and H). In cerebellum, oral delivery of CSP-TTK21 induces histone acetylation, which was revealed by immunofluorescence analysis for H4K12ac (Fig. 7 I-K). The H3K9 acetylation mark was also observed to be upregulated in cerebellum (Fig. 7, L and M) of SCI rats treated with CSP-TTK21 as compared to that with vehicle CSP. Collectively, these results suggest that CSP-TTK21 treatment induces histone acetylation especially in prefrontal cortex and cerebellum of SCI rats which might results in activation/expression of RAGs leading to axonal regeneration and hence functional recovery as observed on motor functions in these rats.

**Fig. 7.**
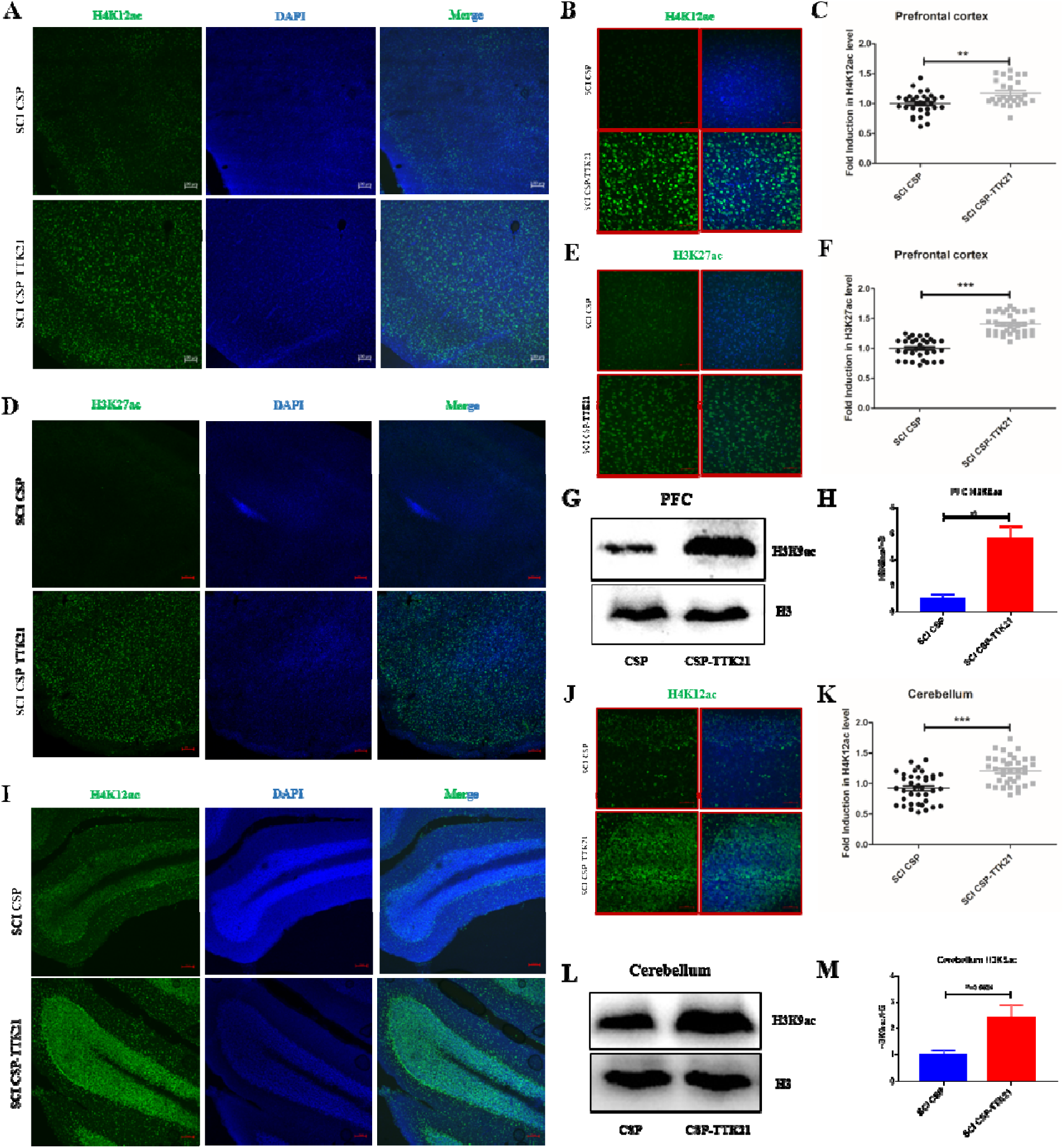
Histone acetylation in prefrontal cortex and cerebellum in CSP-TTK21 treated rats in spinal cord injury. (A) Immunofluorescence image showing increased H4K12ac (green) in prefrontal cortex. (B) Enlarged image at 40X magnification showing the induction of H4K12ac marks upon CSP-TTK21 treatment. Scale bar (A) 100 µm and (B) 50 µm. (C) Quantification of fold induction in H4K12ac levels in prefrontal cortex of SCI CSP-TTK21 rats as compared to SCI CSP rats (5-6 rats/group). (D) Immunofluorescence image showing increased H3K27ac (green) in prefrontal cortex. (E) Enlarged image at 40X magnification showing the induction of H3K27ac marks upon CSP-TTK21 treatment. Scale bar (D) 100 µm and (E) 50 µm. (F) Quantification of fold induction in H3K27ac levels in prefrontal cortex of SCI CSP-TTK21 rats as compared to SCI CSP rats (5-6 rats/group). (G) Immunoblot analysis showing increased H3K9ac marks in prefrontal cortex (H) Quantification of fold induction in H3K9ac of SCI CSP-TTK21 rats as compared to SCI CSP rats (3-4 mice/group). (I) Immunofluorescence image showing increased H4K12ac in cerebellum. (J) Enlarged image at 40X magnification showing the induction of H4K12ac marks upon CSP-TTK21 treatment. Scale bar (I) 100 µm and (J) 50 µm. (K) Quantification of fold induction in H4K12ac levels in cerebellum of SCI CSP-TTK21 rats as compared to SCI CSP rats (5-6 rats/group). (L) Immunoblot analysis showing increased H3K9ac marks in cerebellum (M) Quantification of fold induction in H3K9ac of SCI CSP-TTK21 rats as compared to SCI CSP rats (3-4 mice/group). Student’s t-test was done for significance test, **pLJ0.01, ***pLJ0.001.

### CSP-TTK21 promotes expression of regeneration-associated genes in prefrontal cortex and cerebellum after spinal cord injury

Next, we wanted to establish whether induction of acetylation by oral delivery of CSP-TTK21 observed in prefrontal cortex and cerebellum is correlated with regeneration-associated genes (RAGs) expression after spinal cord injury. We therefore tested whether oral treatment of CSP-TTK21 would be sufficient to induce RAGs in prefrontal cortex and cerebellum. To this end, we selected 11 of the well-known RAGs for our study and checked their expressions by quantitative real-time PCR. We observed that oral delivery of CSP-TTK21 induced expression of 6 RAGs out of the 11 RAGs (Gap43, Klf7, Smad1, Sprr1a, Npy and S100b) in the prefrontal cortex (Fig. 8A) of SCI rats. Additionally, we observed increased expression of 4 key RAGs (Gap43, Smad1, Npy and S100b) in the cerebellum region of CSP-TTK21 treated SCI rats (Fig. 8B).

Importantly, these results suggest that targeted pharmacological activation of CBP/p300 lysine acetyltransferase activity by oral delivery of small molecule activator CSP-TTK21 enables simultaneous induction of multiple RAGs. Notably, a weekly 5 dosage of CSP-TTK21 treatment promotes motor recovery after 5 weeks, a finding perhaps associated with a cascade of gene induction. Gap43 (Growth-associated protein-43) is an axonal phosphoprotein and is believed to be an essential factor in promoting axonal growth. Increased mRNA levels of GAP43 have been correlated with axonal sprouting and enhanced regenerative capacity [28]. We observed increased expression of Gap43 in prefrontal cortex and cerebellum of SCI rats upon treatment of CSP-TTK21, which might be contributing to recovering from injured state. Another RAG whose expression was increased upon CSP-TTK21 treatment was Smad1. Smad1 is an intracellular modulator of BMP signalling in DRG neurons. Previously, it has been shown that selective reactivation of Smad1 promotes regeneration of sensory axons in a rodent SCI model [29]. Beside these, expression of Neuropeptide Y (NPY) was also found to be increased upon CSP-TTK21 treatment. NPY has been implicated in neuroprotection and neurogenesis [30]. S100b is a calcium-binding protein and is primarily expressed in astrocytes. This protein is critical for neuronal survival and promotes neurite extension. S100b level has been observed to be upregulated in serum and in cerebrospinal fluid after spinal cord injury to prime the regeneration [31]. We also observed an increased expression of S100b upon CSP-TTK21 treatment. further supporting the required regeneration for the functional recovery. Collectively, these results suggest that CSP-TTK21 increases expression of RAGs and therefore results in functional recovery as observed on motor functions in these rats.

**Fig. 8.**
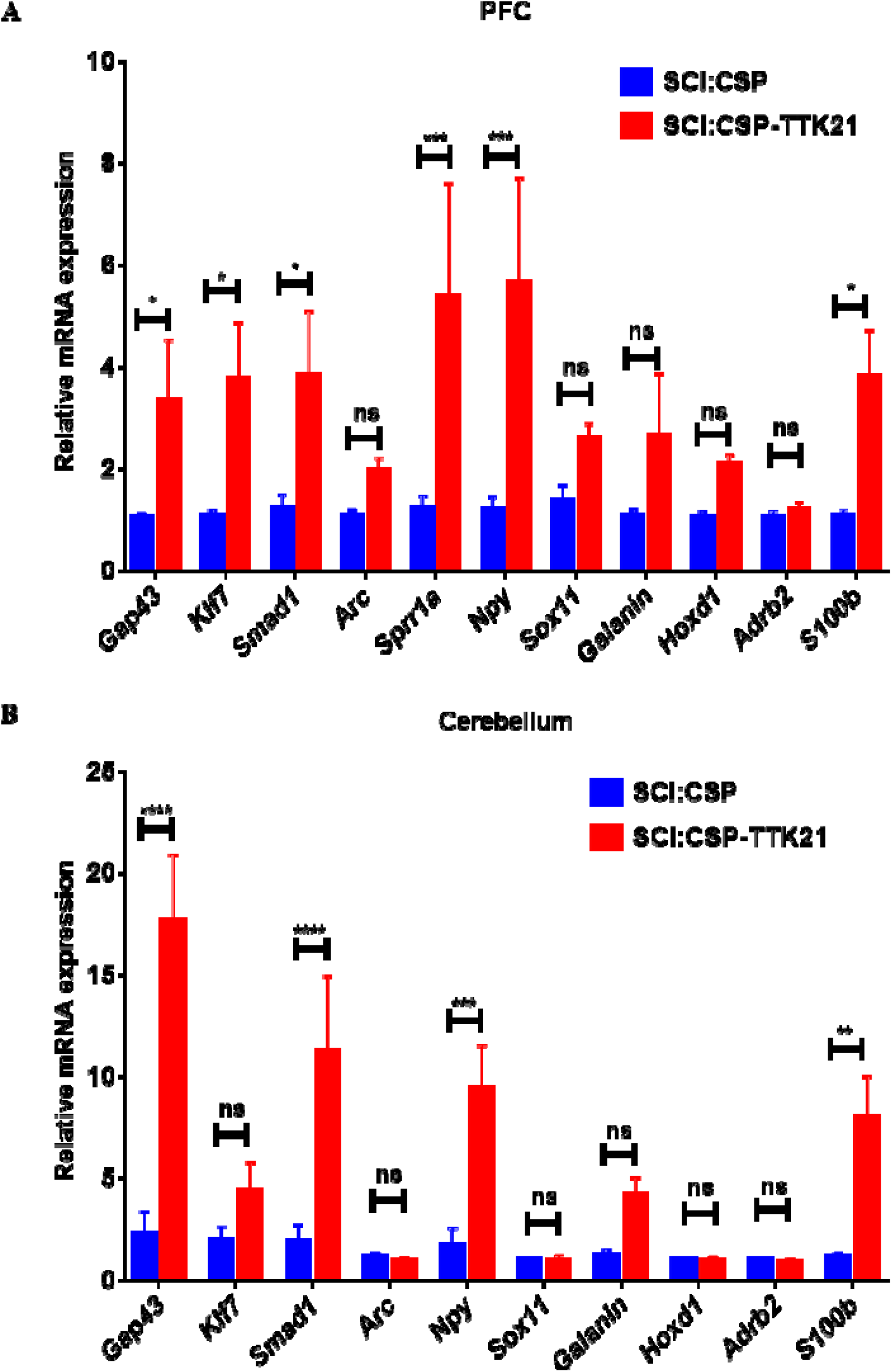
Expression of RAGs in prefrontal cortex and cerebellum in CSP-TTK21 treated rats in spinal cord injury. (A) qRT-PCR results of RAGs in mRNA extracts from prefrontal cortex of SCI CSP-TTK21 rats as compared to SCI CSP rats (3-4 rats/group). (B) qRT-PCR results of RAGs in mRNA extracts from cerebellum of SCI CSP-TTK21 rats as compared to SCI CSP rats (3-4 rats/group). Two-way anova with uncorrected Fisher’s LSD test was performed for statistical analysis, *p LJ 0.05, **p LJ 0.01, ***p LJ 0.001 and ****p LJ 0.0001.

Our research has unveiled promising insights into the potential therapeutic applications of CSP-TTK21. We’ve successfully demonstrated that CSP-TTK21 can be effectively delivered to the brain through oral administration, triggering histone acetylation and promoting long-term potentiation (LTP) within the hippocampal region of wild-type mice. Notably, when we conducted a comparative assessment of CSP-TTK21’s performance between Oral Gavaging and IP administration, we made a remarkable discovery—both methods exhibited nearly identical levels of activity and functionality. This groundbreaking study represents the first direct exploration of CSP-TTK21’s role in synaptic plasticity through the induction of LTP. The implications of these findings are substantial, signifying the immense therapeutic potential of CSP-TTK21 for a wide spectrum of neurological disorders [32], all while highlighting its feasibility for oral administration. Moreover, our investigation has extended beyond the realm of CSP-TTK21 alone. We have also explored the utility of CSP as an oral drug delivery system, particularly for targeting brain-related ailments such as cancer, neurodegenerative diseases, brain tuberculosis, and spinal injuries.

Intriguingly, we extended our study to investigate the effects of orally administering CSP-TTK21 in a rat spinal cord injury model. Our observations revealed that oral administration of CSP-TTK21 not only significantly improved behavioural measures but also concomitantly induced histone acetylation, effectively contributing to the repair of spinal cord injuries. These findings hold immense promise for the development of novel treatments and therapies for spinal cord injuries and various neurological disorders. Collectively, these results establish that oral treatment of the glucose-derived carbon nanosphere (CSP) conjugated small molecule activator of p300/CBP acetyltransferase actively induces acetylation and thereby the axonal growth which promotes spinal injury repair resulting in functional recovery of the injury related paralysis (Fig. 9) [9, 15].

**Figure 9.**
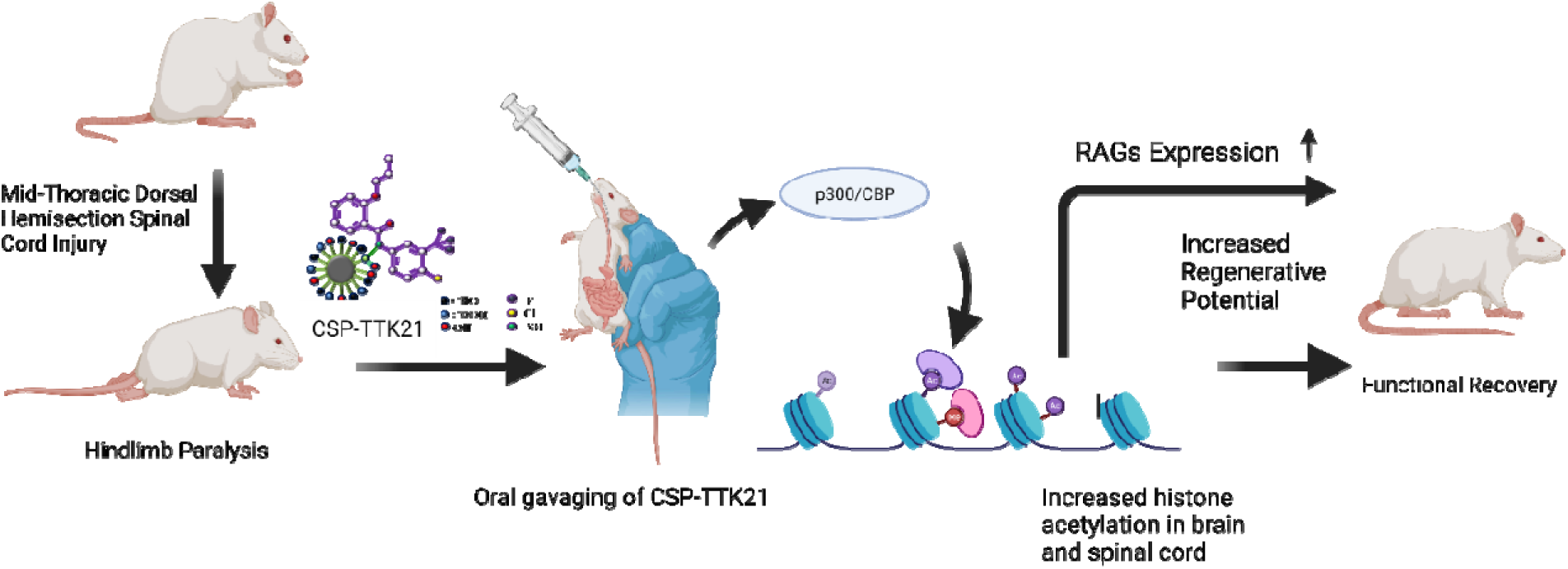
The proposed model for CSP-TTK21 mediated improvement in motor functions after SCI. We propose that oral administration of CSP-TTK21 improves motor activities after SCI by restoring histone acetylation dynamics through activation of CBP/p300 KATs activities resulting in rescue of transcription dynamics, especially by restoring the expression of regenerative associated genes. (Created with BioRender.com)

## MATERIALS AND METHODS

### Animal housing

C57/BL6 mice were procured from The Jackson Laboratory. All animals were kept in standard mouse cages under conventional laboratory conditions (12h/12h light-dark cycle, temperature: 22±2°C, humidity: 55±5%) with *ad libitum* access to food and water. 4-8 weeks old mice were used for each experiment. All the experiments were performed in accordance with Institutional Animal Ethics Committee (IEAC), and the Committee for the Purpose of Control and Supervision of Experiments on Animals (CPCSEA).

Adult male Sprague Dawley (SD) rats weighing 80-100 grams were procured from the National Laboratory Animal Centre (NLAC) of Central Drug Research Institute, Lucknow, India. Rats were kept in a pathogen-free environment at constant temperature (22±2°C) and 12/12 hrs. Light/dark cycle (8:00 a.m. to 8:00 p.m.) with free access to food and water. All animal protocols and experiments are approved by the Institutional Animals Ethics Committee (IAEC) following the guidelines of Committee for the Purpose of Control and Supervision of Experiment on Animals (CPCSEA), which complies with international norms of INSA (Indian National Science Academy).

### Immunofluorescence and Immunohistochemistry for animal tissue

CSP and CSP-TTK21 were administered orally to mice at a dose of 20 mg/kg of body weight and for rats 10 mg/kg of body weight. Mice were killed via cervical dislocation after 84 hour and rats by carbon dioxide euthanasia procedure after conclusion of behaviour experiments. The brain and spinal cord was removed, rinsed with PBS, and then put in 4% paraformaldehyde (PFA) for a 24-hour fixing period at room temperature. Tissues were then maintained at 4°C in 30% sucrose while being prepared for cryosectioning. The tissues were frozen in optimal cutting temperature (OCT-Leica) media for 20 min at -20°C. Using a Cryotome (Leica), thick coronal slices measuring 20 µm for mice were cut through the dorsal hippocampus. The rat brain and spinal cord coronal cryosections were taken at 40 µm. The tissue slices were permeabilized for 15 min in 1X PBS/0.3% Triton X-100. 1X PBS/0.1% Triton X-100/5% horse serum was used for 30 min at 37°C to inhibit nonspecific labelling. Then the sections were separately incubated with the indicated antibodies (rabbit H3K9Ac, H3K14Ac, H3K27ac and H4K12Ac) overnight. Sections were washed thrice with 1X PBS and then incubated with the appropriate secondary antibody. For immunofluorescence, sections were incubated with goat anti-rabbit Alexa fluor 568 dye [Thermo Fischer A-11011] and anti-rabbit Alexa fluor 488 dye [Thermo Fischer A-11070] for 1 h at room temperature, under dark, the nuclei were stained with Hoechst [Invitrogen, #H1399] (1:1000 dilutions) or DAPI (1:1000) for 5 min, followed by 3 washes with 1X PBS/0.1% Triton X-100. Then sections were mounted on slides using 70% glycerol. Fluorescence images were taken through confocal scanning laser microscope LSM510 META and Zen LSM810. For immunohistochemistry, after primary antibody sections were further incubated with anti-rabbit horseradish peroxidase-conjugated antibody (#sc2004; Santa Cruz Biotechnology) for 1 h. After three washes with 1X PBS/0.1% Triton X-100, 0.05% DAB (with 0.04 M Tris, pH 7.5, 0.03% H2O2) was added and sections were mounted with a Roti Histokit II (Roth). Carl Zeiss light microscope was used to capture the images for immunohistochemistry, and Zen black software was used to process the data.

#### Brain Lysates

After animal sacrifice, the brain and spinal cord was swiftly removed and thoroughly washed with ice-cold 1X PBS to remove the blood samples. The prefrontal cortex, hippocampus, cerebellum and spinal cord regions were dissected out and homogenised in lysis buffers (RIPA or Laemmli) containing protease inhibitors and sodium butyrate (NaBu). Homogenised samples were centrifuged for 20 minutes at 12,000rpm at 4°C. The supernatant was collected, and protein content was determined using the Bradford method with Bio-Rad’s protein assay and RC-DC kit. For future usage, the samples were aliquoted and kept at -80°C. For cell lysates, 10^6^ cells were seeded in each well of 6-well plates and grown at 37 °C. At about 80% confluency, cells were exposed to various dosages of CSP (50 µg/ml and 100 µg/ml) and CSP-TTK21 (50, 100, and 150 µg/ml) for 12 hours. Cells were collected and then centrifuged for five minutes at 2000 rpm and 4°C. Following a cooled 1X PBS wash, the cells were centrifuged. The pellet was re-suspended in 1X Laemmli buffer (62.5 mm Tris-HCl pH 6.8, 25% glycerol, and 2% SDS) in 10 times the volume of the produced cell pellet. The cell suspension was homogenised using pipette tips, and the homogenate was then heated to 90°C for 10 minutes and cooled to room temperature (RT) for 10 minutes.

### SDS-PAGE and Western Blotting

Protein samples were placed onto an SDS-PAGE gel and electrophoresed. Small and large molecular weight proteins were transferred to the PVDF membrane at 25 V for 25 minutes and 80 V for 1.5 hours, respectively. Blots were stained with Ponceau/Direct Blue after transfer, followed by a PBS wash. After destaining, the blots were blocked for an hour with 5% skim milk (5% BSA for phospho-proteins), washed once with PBS, and then probed with rabbit primary antibodies against H3 (1:10000), H3K9Ac (1:2000), H3K14Ac (1:2000), for an overnight period at 4°C. Blots were washed three times with PBST (PBS+ Tween 20) before being probed with HRP conjugated secondary antibody (Goat anti-rabbit; 1:10000) for one hour at room temperature. The blots were washed three times with PBST before being developed using the Luminol chemiluminescent technique with ECL western clarity solution in a Syngene/Bio-Rad Gel-documentation system. Images were merged and bands quantified using ImageLab version 5.2.1 and ImageJ software, respectively.

### Preparation of hippocampal slices

Acute brain slices were prepared from C57/BL6 male and female adult mice (>6 weeks). Mice were sacrificed by cervical dislocation, brains were dissected and stored in cold sucrose-based cutting solution (124 mM Sucrose, 3 mM KCl, 24 mM NaHCO_3_, 2 mM CaCl_2_, 1.25 mM NaH_2_PO_4,_ 1 mM MgSO_4_, and 10 mM D-Glucose) bubbled with carbogen (5% CO_2_ and 95% O_2_; Chemix, India). The brain was glued to a vibratome holder, and 350 µm thick horizontal hippocampus slices were sliced in an ice-cold sucrose cutting solution. Slices were stored for an hour in a slice chamber containing artificial cerebrospinal fluid (aCSF: 124 mM NaCl, 3 mM KCl, 24 mM NaHCO_3_, 2 mM CaCl_2_, 1.25 mM NaH_2_PO_4_, 1 mM MgSO_4_, and 10 mM D-Glucose) bubbled with carbogen. Following recovery, the slices were stored at room temperature in the slice holder until further usage.

### Extracellular field recordings

Slices were transferred one at a time to a submersion type recording chamber continuously perfused with carbogenated aCSF and maintained at 34°C with in-line solution heaters. Extracellular field Excitatory Post Synaptic Potentials (fEPSPs) were recorded from the Schaffer-Collateral commissural pathway of the hippocampus. fEPSPs were elicited by electric stimulation of CA1 region of stratum radiatum with a bipolar stimulating electrode (CBARC75, FHC, United States), and fEPSPs were recorded from the CA3 region of the stratum radiatum using low-resistance (3-5 M) glass electrodes filled with aCSF (pH 7.3) which were pulled from borosilicate glass capillaries (ID: 0.6mm, OD: 1.2mm, Harvard Apparatus) using a horizontal micropipette puller (Flaming-Brown P-97, Sutter Instruments Co., USA). A stimulation frequency of 0.1 Hz was employed. Input-output curves were generated by adjusting the stimulus intensity in increments of 20 µA for each sweep. These increments ranged from 0 to 160 µA, with a stimulation duration set within the range of 20 to 40 µs. To assess Paired-pulse Facilitation (PPF) before establishing the baseline, we utilized a series of paired pulses separated by quarter log unit intervals, covering intervals from 10 to 1000 milliseconds. Slices that did not exhibit stable baseline activity for a duration of 15-20 minutes following its acquisition were excluded from the study. To induce long-term potentiation (LTP), a single train of 100Hz stimuli was administered over the course of one second. Subsequently, the resulting activity was closely monitored for a period of 55 to 60 minutes.Signals were amplified using Molecular Devices’ Multiclamp 700B, filtered at 2 kHz, then digitalized at 20–50 KHz using its Digidata 1440A. Data collections were carried out using Molecular Devices’ pClamp 10.2 software, and Clampex 10 software was used for analysis.

### Dorsal hemisection injury

In brief, animals were anesthetized using a mixture solution consisting of 35 mg/kg of ketamine and 8 mg/kg of xylazine based on their body weight. The depth of anesthesia was assessed by manually pinching the hindlimbs; a sufficiently anesthetized animal displayed a breathing response but no limb movement. Next, the procedure involved the following steps:

1. Shaving and triple cleaning of the skin over the thoracic area.
2. Incision of the skin, separation of superficial fat, and dissection of muscle tissue to expose laminae T9-T11.
3. Performing a T9 laminectomy while ensuring the preservation of the spinal cord by removing the duramater.
4. Creating a mid-thoracic dorsal hemisection injury at the T9 level using a 24 G injection needle and microscissors. Confirmation of the complete injury was evident through the animal’s reaction, including leg extension and tail movement.
5. Closure of the detached muscle and skin, followed by the return of the animals to their cages.

Postoperatively, all injured animals received antibiotics (enrofloxacin, 10-20 mg/kg body weight for 7 days) and an analgesic (meloxicam at 2 mg/kg + paracetamol at 60 mg/kg body weight for 7 days). Throughout the experiment, animals underwent daily monitoring for general health, mobility within the cage, wound assessment, swelling, infection, and autophagia of toes as reported previously [33].

Additionally, the animals received oral administration of 0.9% saline (10 ml/kg body weight) twice daily for three days and once daily from day 4 to 7 post-surgery. Bladder expression was conducted twice daily during the first week after surgery and then once daily as needed.

### CSP-TTK21 administration in SCI rats

Animals were randomized and divided into two groups to the treatment with HAT activator (CSP-TTK21) or vehicle control of carbon nanosphere (CSP) (Rats – 10 mg/kg body weight through oral route once per week). Animals received the first oral dose of CSP-TTK21, 4 hours after spinal cord injury to stimulate a clinically feasible delay to treatment and then once a week thereafter for 5 weeks and 9 weeks. No adverse effects were observed following oral administration of CSP-TTK21.

### Open field activity test

To assess locomotor activity, rats were tested in the open-field arena. Animals were individually placed in locomotor boxes to assess horizontal and rearing activity. Prior to starting the experiment, rats were habituated for 10 minutes, and locomotor activity was monitored over a period of 30 minutes. Horizontal activity was calculated in terms of distance travelled over a period of 30 minutes and vertical activity is recorded with the number of rearing counts. Before introducing the next animal into locomotor boxes, the open-field arena was swabbed every time with 10% alcohol to avoid the odour interference from the previous animal. All animal activities were measured using a photoelectric actimeter (Acti-Track System, Panlab).

#### qRT-PCR

Three biological samples of rat brain tissues were used for each group (SCI CSP and SCI CSP-TTK21). On the completion of treatment period, tissues were collected, snap frozen in liquid nitrogen, and kept at -80°C. The cerebellum and prefrontal cortex was removed, and the total RNA was isolated through further processing. The Trizol method (Invitrogen Trizol 15,596-026) was utilized to isolate total RNA from the cerebellum and prefrontal cortex. The tissues were homogenized in 200 μl of Trizol reagent, frozen in liquid nitrogen and kept at a temperature of - 80°C. Samples were thawed on ice and then centrifuged at 13,000 rpm for RNA isolation. The supernatant underwent RNA extraction using phenol:chloroform. RNA was precipitated at room temperature (15 min) using iso-propanol and RNAse-free glycogen. After a single 75% ethanol wash, the pellet was allowed to air dry for five to ten minutes before being resuspended in DEPC-treated/RNAse-free water. DNAse treatment (30 min at 37°C) was applied to isolated RNA samples, and then they were re-precipitated and resuspended (10 min at 55°C). Following the manufacturer’s recommendations, two μg of RNA were utilized for cDNA synthesis using MMLV-reverse transcriptase and oligo-dT (Sigma: M 1302 and O4387, respectively). A CFX96 Real-Time PCR Detection System with a C1000 Thermal Cycler (Bio-Rad) was used to perform qPCR using 2× Takara TB Green Mastermix (ABI), and the respective specific primers are shown in Table 1. The data were analyzed using Bio-Rad CFX Maestro Software version 4.1. Fold changes were calculated using the formula: 2(-[Ct Test-Ct Control]). Wherever it applied, Gapdh was regarded as a housekeeping gene.

**TABLE 1.**
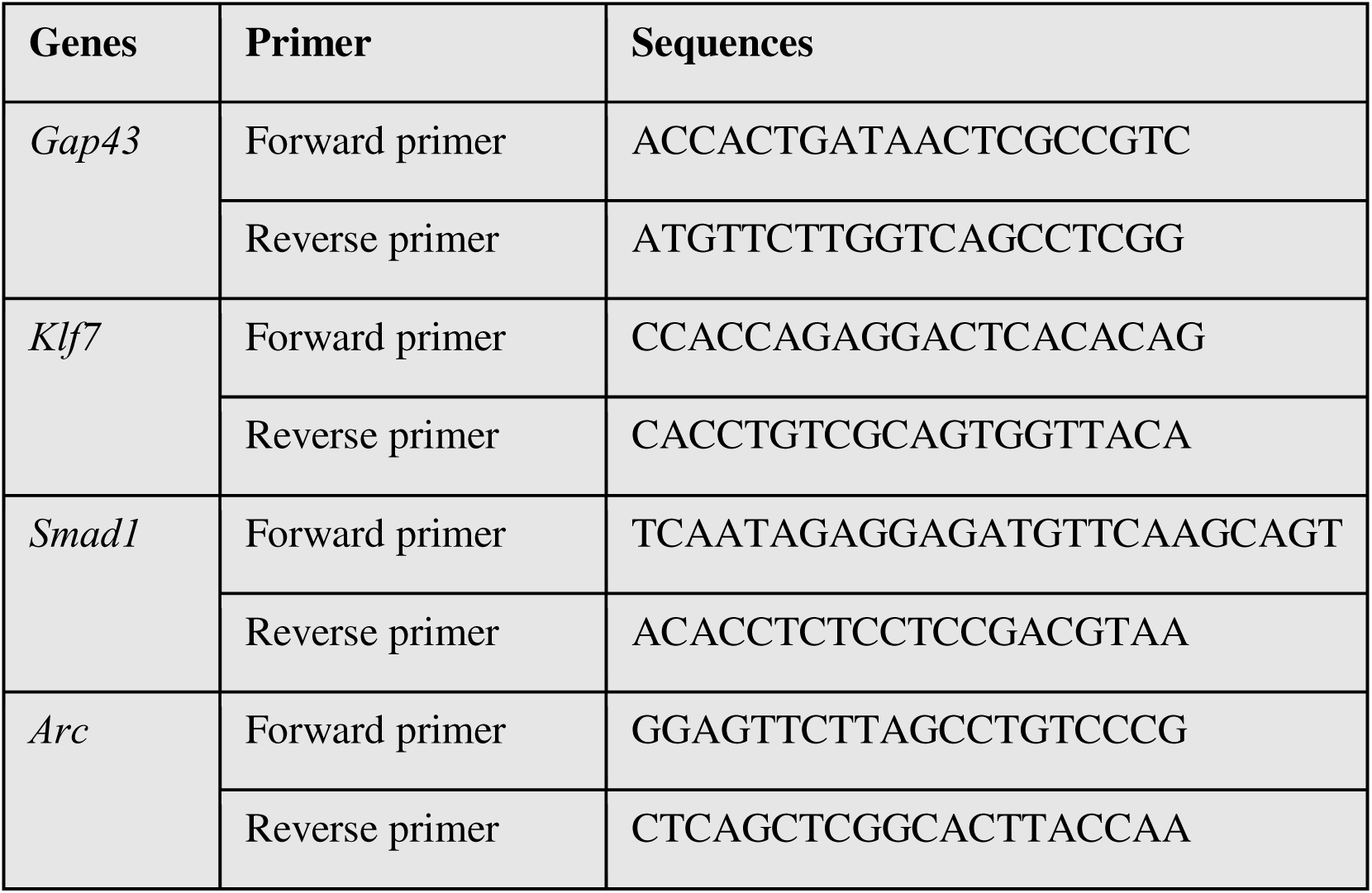
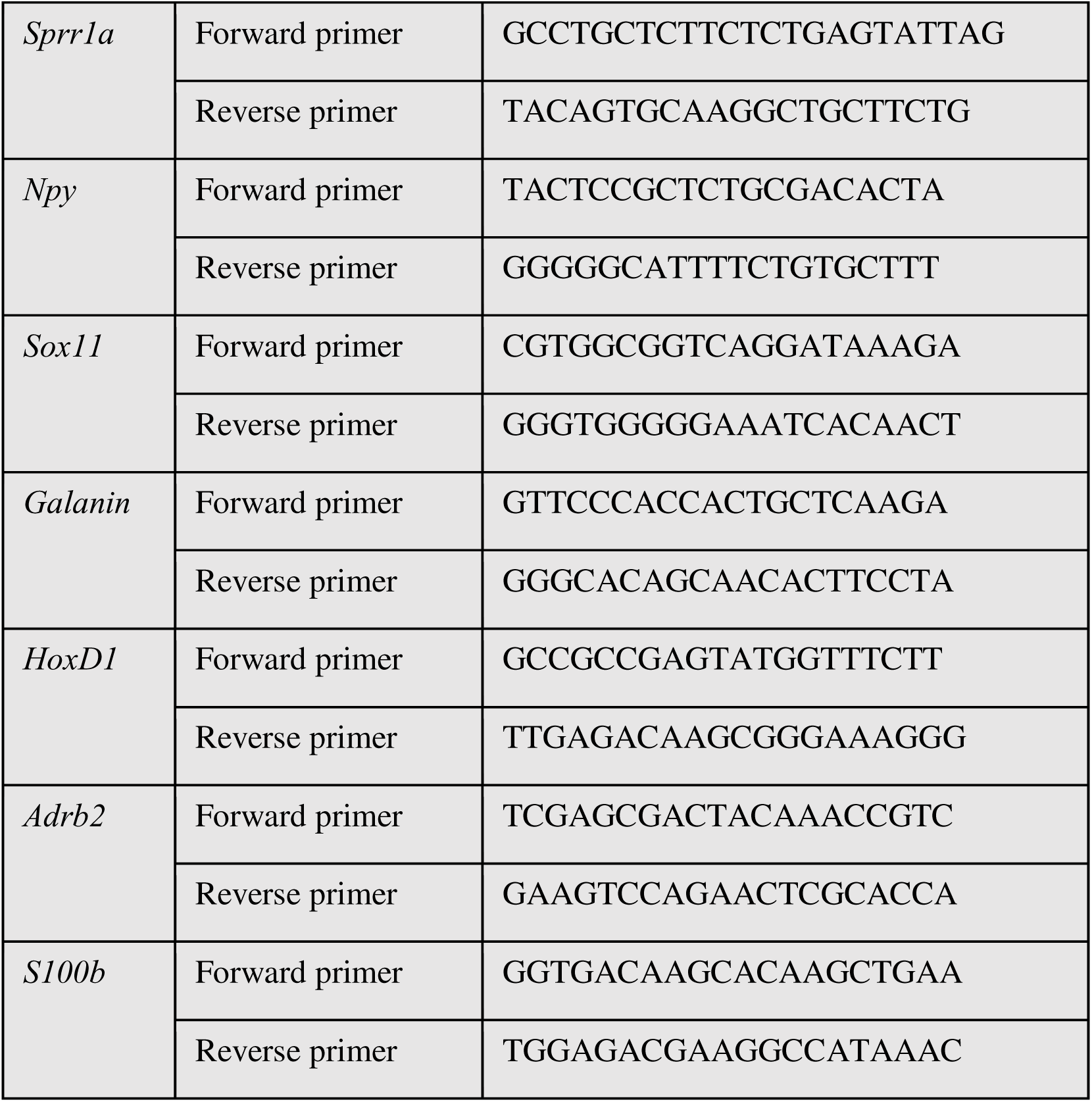
Primer sequences used for qRT-PCR experiments.

#### Statistical analysis and significance test

The GraphPad Prism 7 software (California, USA) was used to conduct all statistical analyses. Data from three or more individual experiments, as indicated in the figure legends, were expressed as mean SD. The statistical significance levels were calculated using the two-tailed unpaired Student’s t-test or the one-way ANOVA with Dunnett’s/Bonferroni’s multiple comparison.

## Acknowledgment

This work was supported by the Jawaharlal Nehru Center for Advanced Scientific Research (JNCASR), CSIR-Central Drug Research Institute, Lucknow, J C Bose Fellowship, Department of Science and Technology (DST), India (grant no. SR/S2/JCB-28/2010) to TKK, AKS is supported by the JNCASR Research Fellowship, and AR is supported by the University Grants Commission (UGC) for predoctoral fellowship.

